# Adding carbon fiber to shoe soles does not improve running economy: a muscle-level explanation

**DOI:** 10.1101/2020.02.28.969584

**Authors:** Owen N. Beck, Pawel R. Golyski, Gregory S. Sawicki

**Affiliations:** George W. Woodruff School of Mechanical Engineering, Georgia Institute of Technology, Atlanta, GA; School of Biological Sciences, Georgia Institute of Technology, Atlanta, GA; Parker H. Petit Institute for Bioengineering and Biosciences, Georgia Institute of Technology, Atlanta, GA

## Abstract

In an attempt to improve their distance-running performance, many athletes race with carbon fiber plates embedded in their shoe soles. Accordingly, we sought to establish whether, and if so how, adding carbon fiber plates to shoes soles reduces athlete aerobic energy expenditure during running (improves running economy). We tested 15 athletes during running at 3.5 m/s in four footwear conditions that varied in shoe sole carbon fiber plate bending stiffness. For each condition, we quantified athlete aerobic energy expenditure and performed biomechanics analyses, which included the use of ultrasound imaging to examine soleus muscle dynamics *in vivo*. Overall, increased footwear bending stiffness lengthened ground contact time (p=0.048), but did not affect ankle (p≥0.060), knee (p≥0.128), or hip (p≥0.076) joint angles or moments. Additionally, increased footwear bending stiffness did not affect muscle activity (all seven measured leg muscles (p≥0.146)), stride averaged active soleus volume, (p=0.068) or aerobic power (p=0.458) during running. Hence, footwear bending stiffness does not appear to alter the volume of aerobic energy consuming muscle in the soleus, or any other leg muscle, during running. Therefore, adding carbon fiber plates to shoe soles slightly alters whole-body and calf muscle biomechanics but does not improve running economy.

## Introduction

In competitive athletics, marginal differences distinguish champions from their competitors. For instance, if any of the top-five 2016 Olympic women’s marathon finishers ran 0.51% faster, they would have been crowned Olympic champion. Such miniscule differences highlight the importance for athletes to optimize all factors that influence race performance. One way to further optimize athletic performance is to don the best footwear. Using footwear that reduces athlete aerobic energy expenditure at a given running speed (improves their running economy) can augment distance-running performance by decreasing their relative aerobic intensity.^1–3^ An established method of improving footwear to augment athlete distance-running performance is to reduce its mass.^1, 2, 4, 5^ Based on literature values, if an aforementioned Olympic marathoner re-raced in shoes that were 100 g less than their original footwear, they would have expended aerobic energy at an ∼0.8% slower rate,^5^ run the marathon ∼0.56% faster,^6^ and taken the gold medal back to their home country.

A longstanding footwear technology that has recently polarized the running community is the use of carbon fiber plates in shoe soles.^7^ Despite the rampant use of carbon fiber plates in athletics,^8–10^ many seek to regulate the use of these plates in distance-running footwear based on the notion that they provide wearers an ‘unfair advantage’ over competitors without such technology.^11^ These views persist even though it is inconclusive whether adding carbon fiber to shoe soles improves running economy.^12–16^ To date, two studies have reported that adding optimally stiff carbon fiber plates to shoe soles improves running economy by 0.8^12^ and 1.1%^13^, while another three studies reported that adding carbon fiber plates to shoe soles does not affect running economy.^14–16^

Moreover, neither study that improved athlete running economy by adding carbon fiber plates to their shoes measured a physiologically-relevant link between the footwear-altered biomechanics and aerobic energy expenditure.^12, 13^ Namely, the first study did not identify a mechanism^12^ while the second suggested that adding carbon fiber plates to shoe soles improves running economy by altering a biomechanical parameter that may not affect metabolism.^13^ Specifically, the second study reported that adding carbon fiber plates to shoe soles improves running economy by decreasing the summed angular impulse (integral of torque with respect to time) during push-off across the leg-joints.^13^ However, angular impulse may not characterize aerobic energy expenditure because generating a given impulse via longer durations (and decreased limb-joint torque) is typically associated with muscle contractions that expend less metabolic energy.^17–19^ Consequently, it remains uncertain whether adding carbon fiber plates to shoe soles improves running economy, and if so how – we need a muscle-level explanation.

After all, muscle contractions dominate whole-body aerobic energy expenditure during locomotion.^20^ Unfortunately, no study has assessed muscle dynamics from athletes running with shoes that have carbon fiber soles. Based on lower limb-joint analyses, which do not necessarily reflect muscle dynamics,^21, 22^ metatarsophalangeal- and ankle-joint dynamics are more affected during running with the addition of carbon fiber plates to shoe soles than knee- and hip-joints.^13, 14, 23–25^ Further, intrinsic foot muscles do not directly affect running economy,^26^ indicating that altered plantar flexor dynamics during running with versus without carbon fiber plates added to shoe soles may help explain changes in running economy.

Theoretically, how does the addition of carbon fiber plates to shoe soles affect athlete plantar flexor dynamics during running? Adding carbon fiber plates to shoe soles increases the footwear’s 3-point bending stiffness^12, 13, 15, 23, 24^ and typically shifts the athlete’s center of pressure more anterior along the foot during ground contact.^23, 24, 27^ These altered footwear biomechanics theoretically increase plantar flexor force generation and decrease the respective muscle fiber’s operating lengths, such changes require more metabolic energy expenditure per contraction.^28, 29^ Additionally, running with stiffer footwear may decrease plantar flexor shortening velocity, reducing muscle metabolic energy expenditure per contraction.^14^ For example, increasing footwear bending stiffness may yield greater plantar flexor force generation because the anterior shift of the athlete’s center of pressure yields a longer moment arm between the ground reaction force (*F_GRF_*) and the ankle-joint center (*R_GRF_*).^13, 23^ Consequently, this longer moment arm leads to a greater GRF-induced ankle-joint flexion moment.^12, 13, 23, 27^ To prevent the ankle-joint from collapsing, plantar flexor muscle-tendons (MTs) generate force (*F_MTs_*) and apply an opposite moment about the joint.

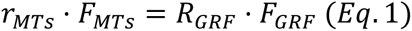

The moment arm between the plantar flexor MTs and ankle-joint center is indicated by *r_MT_*.^30^ When athletes run with carbon fiber plates added to their shoe soles, they often exhibit generally similar GRF magnitudes^13, 15^ and longer plantar flexor MT moment arms^13, 23^ compared to running in conventional footwear. To counteract the greater GRF-ankle flexion moment from adding carbon fiber plates to shoe soles, plantar flexor MT forces increase (Eq. 1). We considered plantar flexor MT force, its physiological cross-sectional area relative to agonist muscles 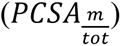,^30^ and pennation angle (*θ_M_*) to estimate how footwear bending stiffness affects plantar flexor muscle fascicle force (*F_M_*).

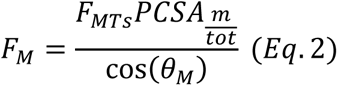

Due to anatomical constraints, the respective MT force increase is likely from greater muscle fascicle force generation; requiring more metabolic energy per contraction.^31^

Adding carbon fiber plates to footwear may also cause plantar flexors to operate at relatively shorter lengths; incurring less economical muscle contractions.^28, 29, 32^ That is because running in footwear that have carbon fiber plates versus without elicits similar leg-joint angles^12, 13^ and MT lengths (*L_MT_*).^33^ Hence, deeming that muscle pennation changes are relatively small, increased MT force may further stretch spring-like tendons (tendon length: *L_T_*) and yield shorter in-series operating muscles (*L_M_*).

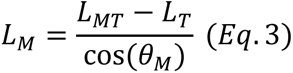

Lastly, adding carbon fiber plates to shoe soles may decrease plantar flexor muscle fascicle shortening velocity during ground contact,^14, 27^ eliciting more economical muscle contractions.^28, 29^ Absent of meaningful changes in ankle-joint mechanical power (*P_ank_*) and plantar flexor MT moment arms (*r_MTs_*), increasing plantar flexor MTs force (*F_MTs_*) decreases ankle-joint angular velocity (*ω_ank_*).^14^

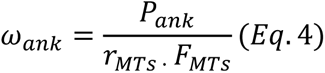

In turn, decreased ankle-joint angular velocity may translate to slower MTs and muscle fascicle shortening velocities.

Perhaps, adding carbon fiber to shoe soles can optimize the trade-off between active muscle force (*P_act_*), force-length (*FL*) and force-velocity (*FV*) relationships to minimize the active volume of plantar flexor muscle^34^ (*V_act_*) (Eq. 5) and aerobic energy expenditure during running.^12, 13^ *σ* is stress and *lm* is fascicle length.

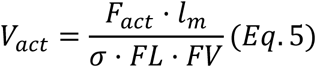

Conceptually, active muscle volume is the amount of muscle that has active actin-myosin cross-bridges, which cyclically split adenosine triphosphate (ATP).^34^ Hence, while considering the rate of metabolic energy expenditure per unit of active muscle volume (*Ṗ_ρ_*), decreasing active muscle volume would turn reduce metabolic energy expenditure (*Ė_met_*).

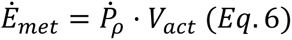

The purpose of this study was to reveal if and how adding carbon fiber plates to shoe soles alters running biomechanics and economy. In particular, we sought to investigate how footwear 3-point bending stiffness affects muscle dynamics and running economy. Based on the reported interactions between adding carbon fiber plates to shoe soles, footwear 3-point bending stiffness,^12–15, 23, 24, 27^ and ankle-joint dynamics,^13, 14, 23, 27^ we hypothesized that running with shoes that have stiffer carbon fiber plates would increase soleus (plantar flexor muscle) force generation while decreasing soleus operating length and shortening velocity during the ground contact. We also hypothesized that an optimal footwear bending stiffness would minimize active soleus volume and aerobic energy expenditure. To test our hypotheses, we quantified ground reaction forces, stride kinematics, limb-joint biomechanics, soleus dynamics, muscle activation patterns, and aerobic energy expenditure from 15 athletes running at 3.5 m/s using four separate footwear conditions that spanned a 6.4-fold difference in bending stiffness (Table 1).

**Table 1.** Participant characteristics. Four and eleven participants initiated ground contact with a mid/forefoot strike (M/FFS) and heel strike (HS), respectively. All participants maintained the same foot strike pattern across footwear conditions.

| Participant | Age<br>(yrs) | Height<br>(m) | Mass<br>(kg) | Leg<br>length<br>(m) | US<br>men's<br>shoe<br>size | Initial<br>Foot<br>Strike | Standing<br>aerobic<br>power<br>(W/kg) |
| --- | --- | --- | --- | --- | --- | --- | --- |
| 1 | 20 | 1.65 | 57.0 | 0.89 | 9 | HS | 1.37 |
| 2 | 27 | 1.73 | 65.6 | 0.91 | 10 | M/FFS | 1.21 |
| 3 | 19 | 1.77 | 60.0 | 0.91 | 9 | HS | 1.98 |
| 4 | 27 | 1.88 | 66.4 | 0.97 | 12 | M/FFS | 1.44 |
| 5 | 27 | 1.70 | 58.9 | 0.83 | 10 | HS | 1.42 |
| 6 | 23 | 1.80 | 72.8 | 0.97 | 10 | HS | 1.70 |
| 7 | 19 | 1.76 | 71.5 | 0.91 | 10 | HS | 1.67 |
| 8 | 24 | 1.78 | 71.3 | 0.93 | 10 | HS | 1.58 |
| 9 | 20 | 1.80 | 66.5 | 0.95 | 10 | HS | 1.72 |
| 10 | 28 | 1.80 | 73.2 | 0.98 | 9 | M/FFS | 1.76 |
| 11 | 28 | 1.74 | 74.5 | 0.83 | 10 | HS | 1.32 |
| 12 | 28 | 1.89 | 75.4 | 0.95 | 12 | M/FFS | 1.21 |
| 13 | 42 | 1.74 | 73.6 | 0.90 | 11 | HS | 1.04 |
| 14 | 26 | 1.79 | 62.2 | 0.90 | 10 | HS | 1.38 |
| 15 | 23 | 1.78 | 65.2 | 0.89 | 9 | HS | 1.28 |
| Average<br>± SD | 25.4<br>± 5.7 | 1.77<br>± 0.06 | 67.6<br>± 6.1 | 0.91<br>± 0.05 | 10.1<br>± 1.0 | 4 M/FFS<br>11 HS | 1.47<br>± 0.26 |

## Results

### Footwear Conditions

Each athlete ran in the Adidas Adizero Adios BOOST 2 running shoes (Adidas) without carbon fiber plates, as well as in the Adidas with 0.8, 1.6, and 3.2 mm thick carbon fiber plates. The average ± SD 3-point bending stiffness for the Adidas was 13.0 ± 1.0 kN/m, and adding 0.8, 1.6, and 3.2 mm thick carbon fiber plates to the shoes soles increased the average ± SD footwear 3-point bending stiffness to 31.0 ± 1.5, 43.1 ± 1.6, and 84.1 ± 1.1 kN/m, respectively. Further, the slope of each footwear-condition’s 3-point bending force-displacement profile was well-characterized by a linear function (average ± SD; Adidas R^2^: 0.97 ± 0.02; Adidas plus in-soles: R^2^: 0.99 ± 0.01).

### Limb-joint Dynamics

Footwear bending stiffness did not affect hip, knee, or ankle angles or moments (Fig. 1). Specifically, footwear bending stiffness was not associated with average, minimum, or maximum ankle (all p≥0.121) (Fig. 1e and Fig. 2g,h), knee (all p≥0.128) (Fig. 1c), or hip (all p≥0.076) angle (Fig. 1a). Similarly, footwear bending stiffness did not affect average or maximum ankle (both p≥0.060) (Fig. 1f), knee (both p≥0.239) (Fig. 1d), or hip (both p≥0.112) (Fig. 1b) moments.

**Figure 1.**
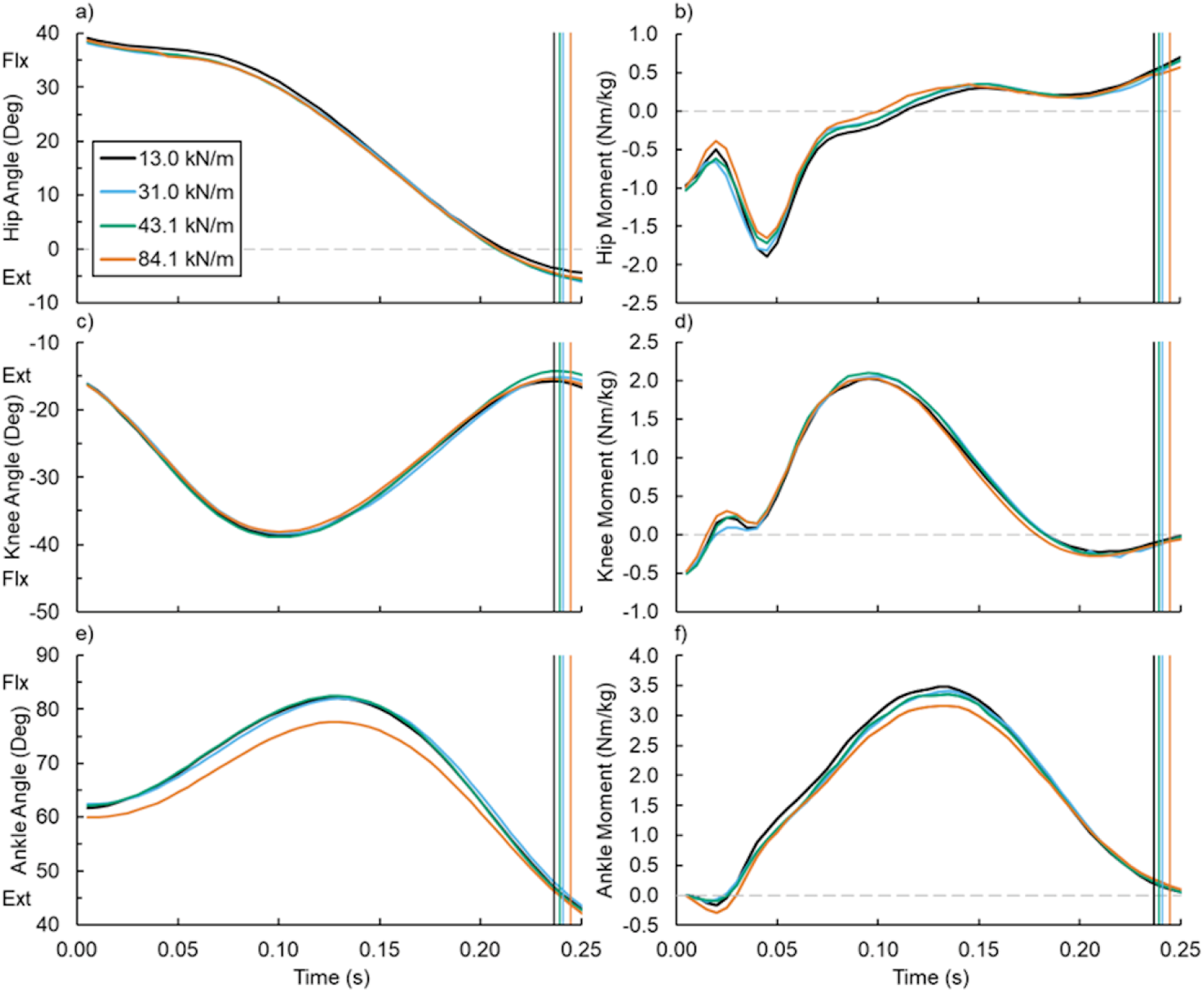
Average (a,b) hip, (c,d) knee, and (e,f) ankle angle and net moment versus time during running with footwear of varied 3-point bending stiffness: 13.0 (black), 31.0 (blue), 43.1 (green), and 84.1 kN/m (orange). Vertical lines indicate the average end of ground contact for the respective footwear condition. Flexion (Flx) and Extension (Ext).

**Figure 2.**
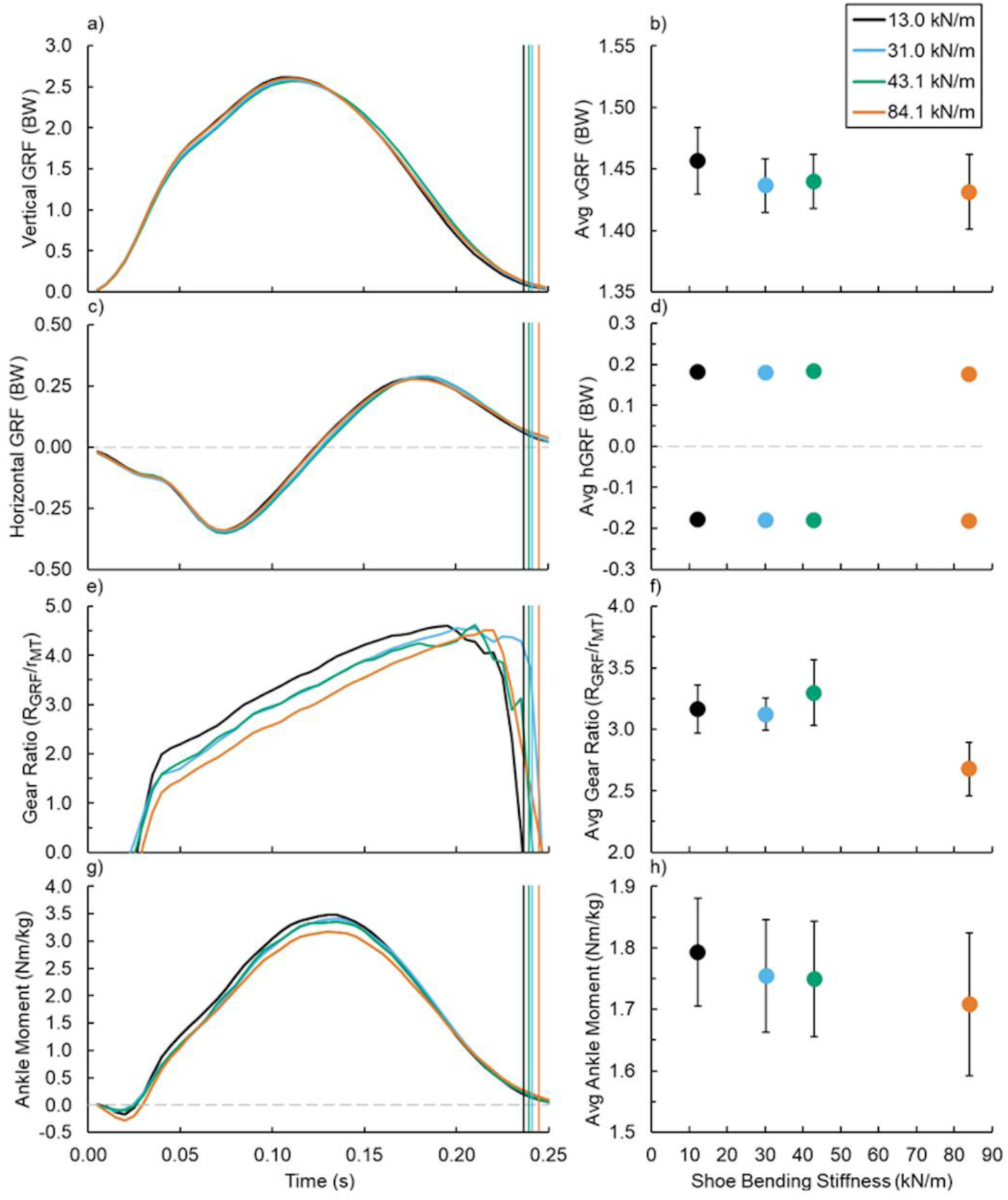
(Left) Average (a,b) vertical and (c,d) horizontal ground reaction force (GRF), (e,f) soleus muscle tendon (MT) gear ratio, and (g,h) net ankle moment versus time and (right) footwear 3-point bending stiffness (right): 13.0 (black), 31.0 (blue), 43.1 (green), and 84.1 kN/m (orange). Vertical lines indicate the average end of ground contact for the respective condition and error bars indicate SE when visible.

### Stride Kinematics and Ground Reaction Forces

Increased footwear bending stiffness was associated with longer ground contact time (p=0.048), but not step time (p=0.956). Regarding GRFs, neither stance average vertical (p=0.209) (Fig. 2a,b), braking (p=0.441) (Fig. 2c,d), nor propulsive (p=0.133) (Fig. 2c,d) GRF differed across footwear bending stiffness conditions. Additionally, footwear bending stiffness did not affect the fraction of vertical (p=0.881) or horizontal (p=0.816) GRF exhibited during the first half of ground contact.

### Muscle-tendon Dynamics

Footwear bending stiffness did not affect soleus muscle-tendon (MT) dynamics (Fig. 3). Neither average soleus MT force (p=0.080) (Fig. 3a,b), length (p=0.150) (Fig. 3c,d), nor velocity (p=0.719) (Fig. 3e,f) during ground contact changed with altered footwear bending stiffness. Additionally, the ratio of the GRF versus soleus MT moment arms to the ankle-joint center (gear ratio) was not affected by footwear bending stiffness (average and maximum gear ratio p=0.371 and p=0.752, respectively) (Fig. 2e,f).

**Figure 3.**
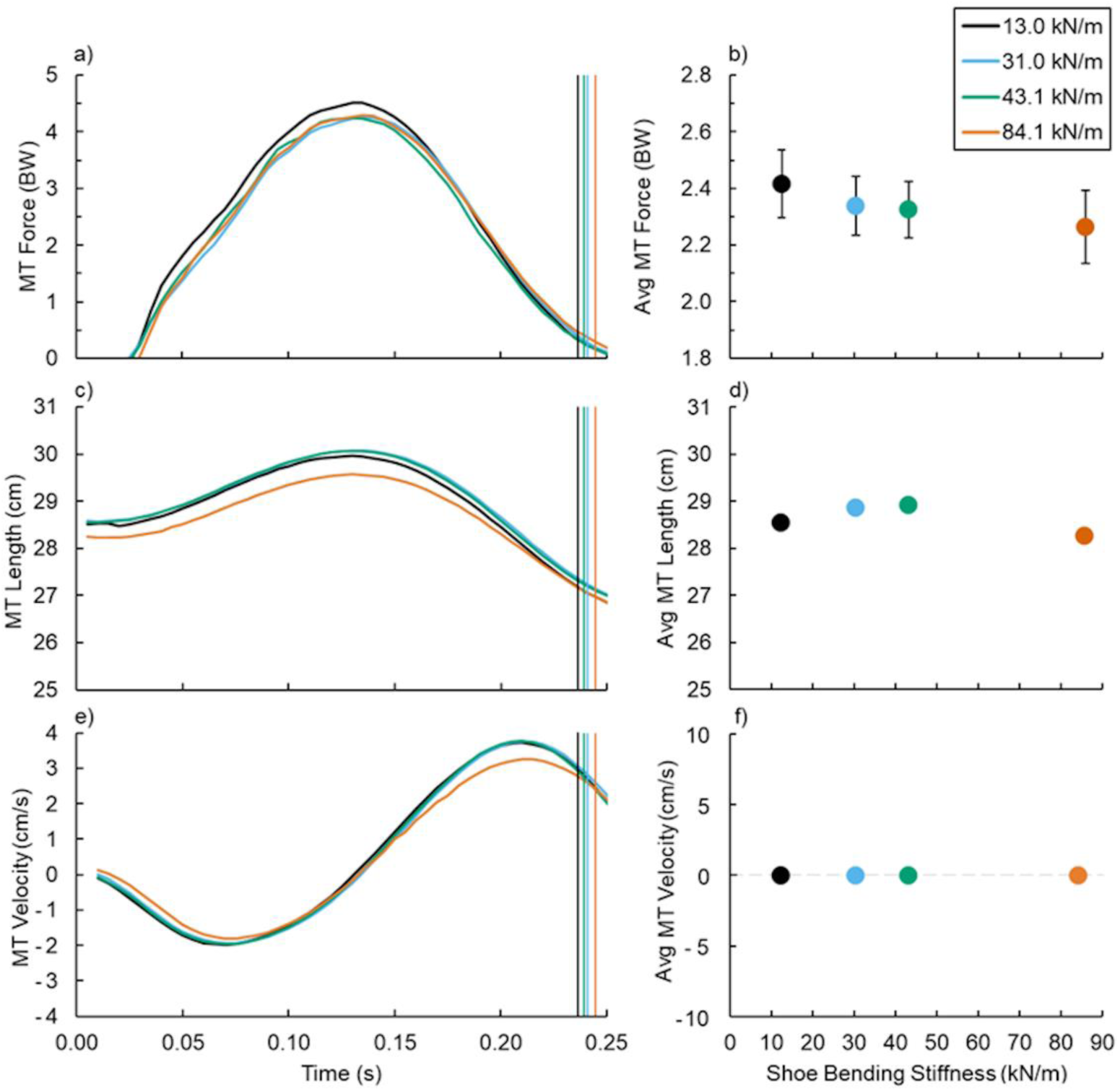
(Left) Average (a,b) soleus muscle-tendon (MT) force, (c,d) length, and (e,f) velocity versus time and (right) footwear 3-point bending stiffness (right): 13.0 (black), 31.0 (blue), 43.1 (green), and 84.1 kN/m (orange). Vertical lines indicate the average end of ground contact for the respective footwear condition and error bars indicate SE when visible.

### Soleus Dynamics

Footwear bending stiffness did not influence average or maximum soleus fascicle pennation angle (both p≥0.476) (Fig. 4a,b), force (both p≥0.115) (Fig. 4c,d and Fig. 5b), velocity (both p≥0.224) (Fig. 4g,h and Fig. 5c), or length (p≥0.286) (Fig. 4e,f and Fig. 5a). Alternatively, footwear bending stiffness did affect the average (p=0.026) and maximum (p=0.014) (Fig. 7b) activated soleus volume during ground contact (Fig. 5d), but did not affect average activated soleus volume per stride (p=0.068) (Fig. 7b & Supplementary Fig. 1a-o).

**Figure 4.**
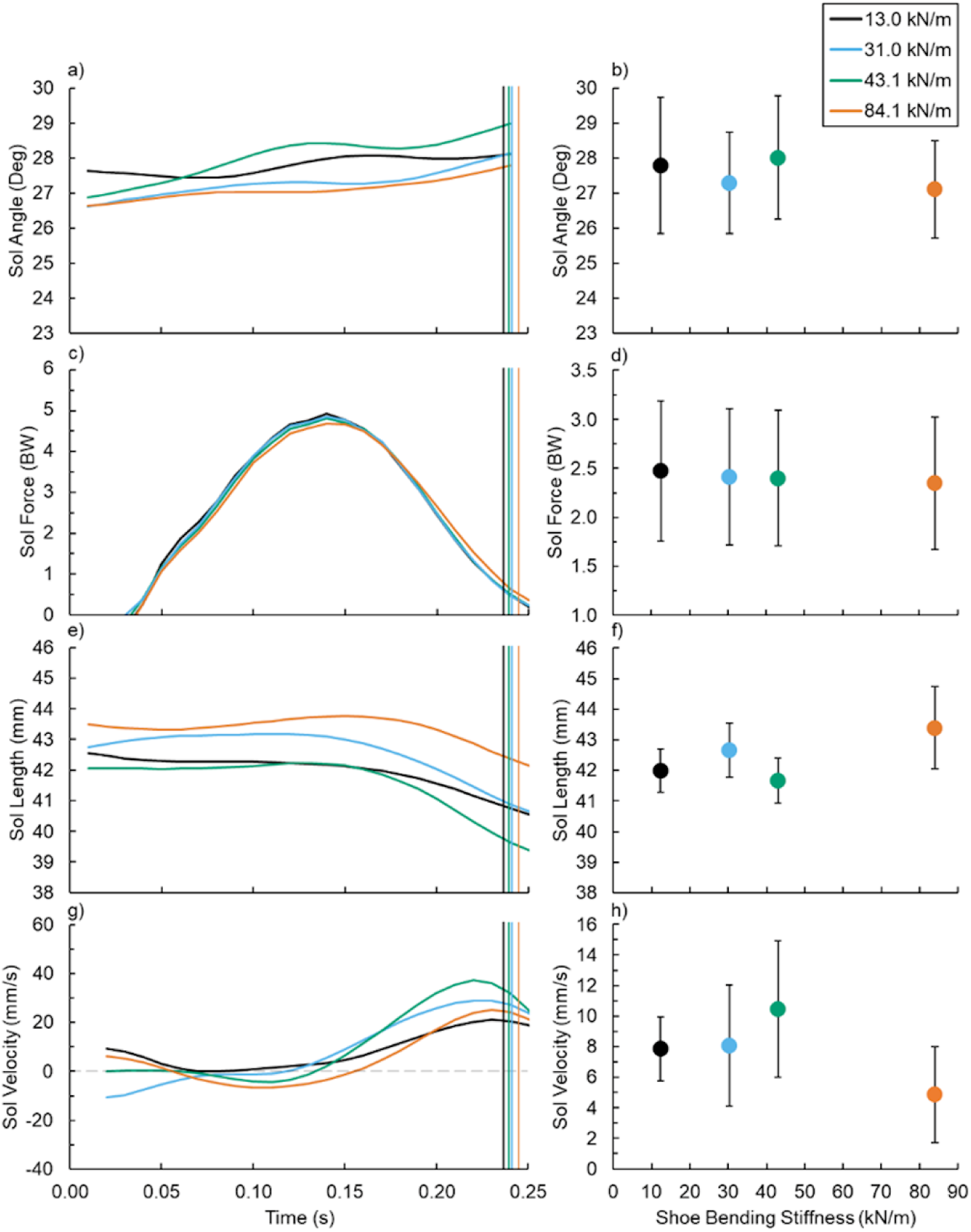
(Left) Average (a,b) soleus (Sol) fascicle angle, (c,d) force, (e,f) length, and (g,h) velocity versus time and (right) footwear 3-point bending stiffness (right): 13.0 (black), 31.0 (blue), 43.1 (green), and 84.1 kN/m (orange). Vertical lines indicate the average end of ground contact for the respective footwear condition and error bars indicate SE.

**Figure 5.**
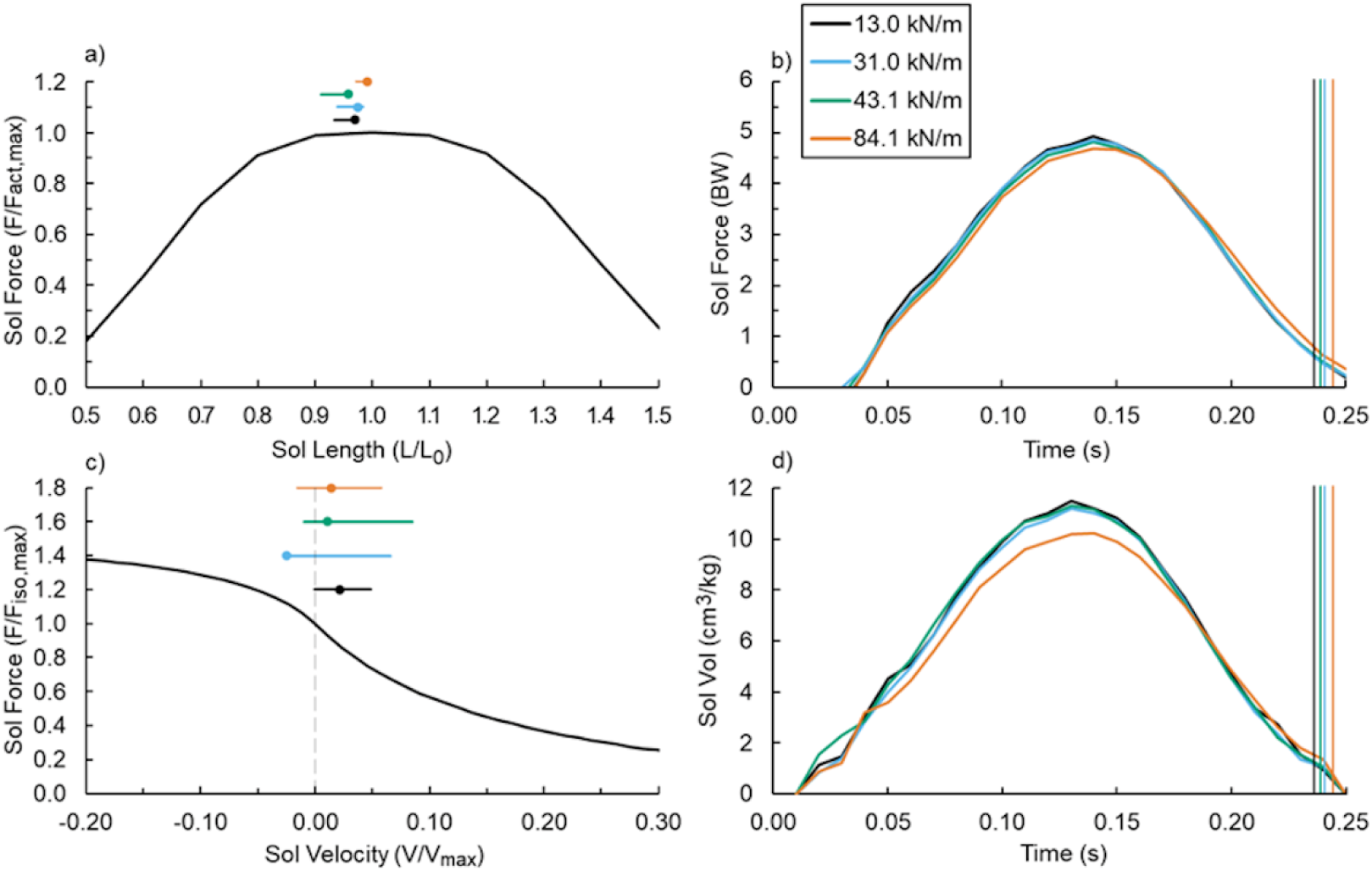
Estimated (a) Soleus (Sol) force-length and (c) force-velocity relationships during ground contact. The marker indicates soleus initial ground contact and the horizontal line indicates soleus operating range during ground contact. (b) Sol force and (d) volume (Vol) throughout ground contact and vertical lines indicate the average end of ground contact.

### Muscle Activation

Footwear bending stiffness did not affect stance- or stride-averaged activation of any measured muscle: soleus (both p≥0.315) (Fig. 6a), medial gastrocnemius (both p≥0.538) (Fig. 6b), tibialis anterior (both p≥0.445) (Fig. 6c), biceps femoris (both p≥0.190) (Fig. 6d), vastus medialis (both p≥0.146) (Fig. 6e), gluteus maximus (both p≥0.603) (Fig. 6f), or rectus femoris (both p≥0.406) (Fig. 6g) (Table 2).

**Figure 6.**
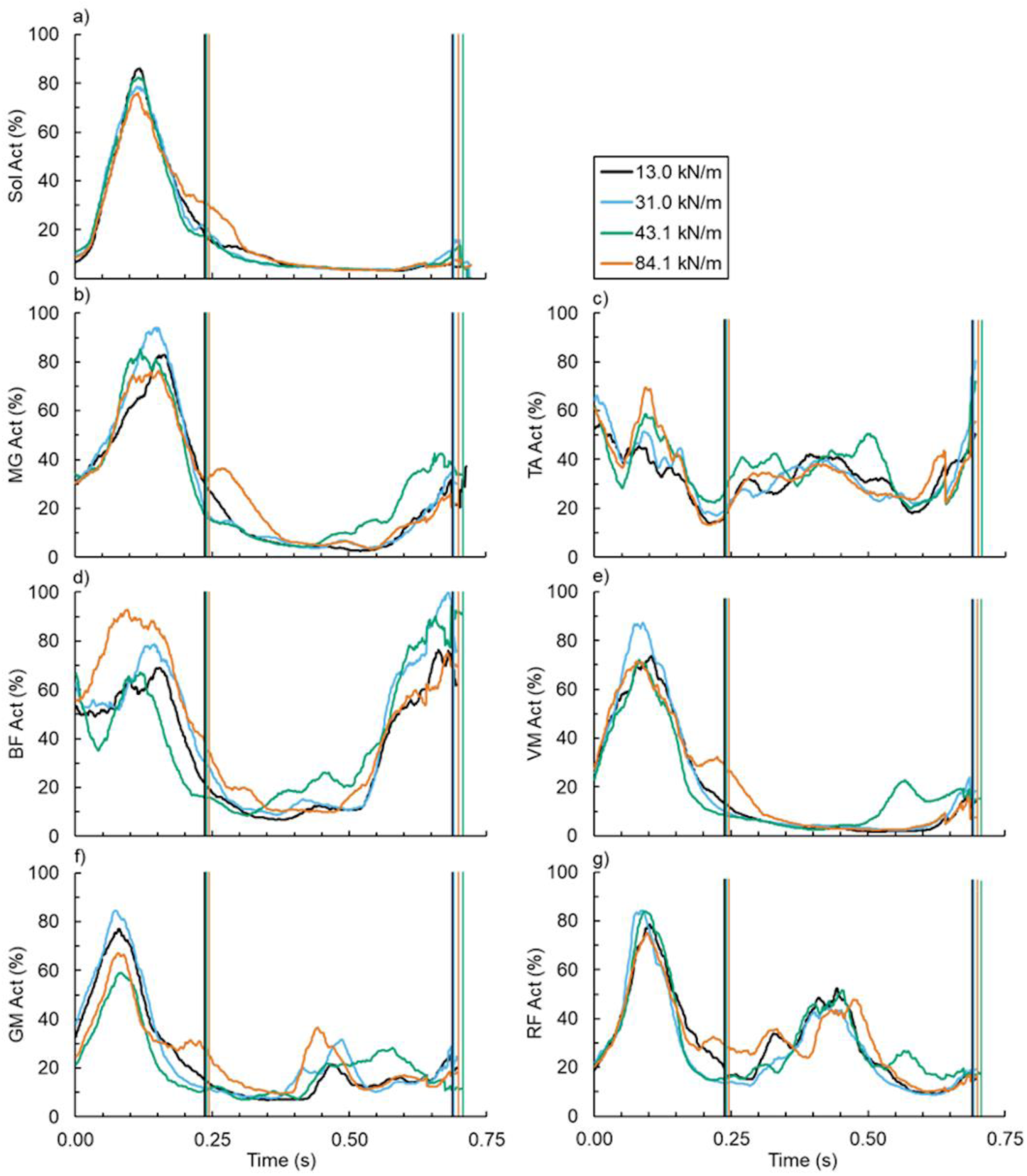
Average (a) soleus (Sol), (b) medial gastrocnemius (MG), (c) tibialis anterior (TA), (d) biceps femoris (BF), (e) vastus medialis (VM), (f) gluteus maximus (GM), and (g) rectus femoris (RF) versus time (left) across footwear 3-point bending stiffness: 13.0 (black), 31.0 (blue), 43.1 (green), and 84.1 kN/m (orange). Vertical lines indicate the average end of ground contact and stride for the respective footwear condition.

**Table 2.**
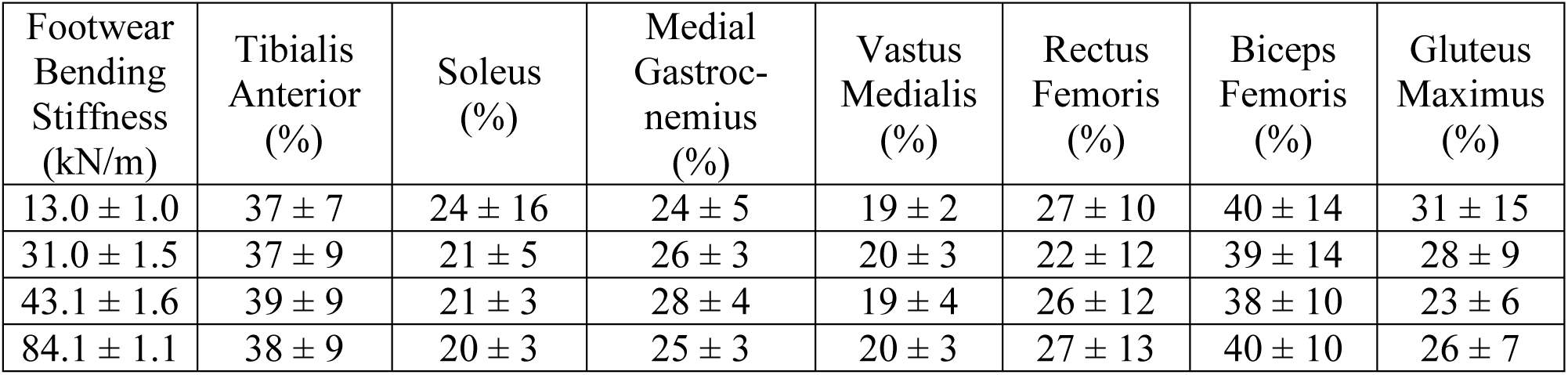
Stride averaged normalized muscle activation (± SD) normalized to the respective muscle’s average maximum value during running with the Adidas (13.0 kN/m) footwear condition.

| Footwear<br>Bending<br>Stiffness<br>(kN/m) | Tibialis<br>Anterior<br>(%) | Soleus<br>(%) | Medial<br>Gastroc-<br>nemius<br>(%) | Vastus<br>Medialis<br>(%) | Rectus<br>Femoris<br>(%) | Biceps<br>Femoris<br>(%) | Gluteus<br>Maximus<br>(%) |
| --- | --- | --- | --- | --- | --- | --- | --- |
| 13.0 ± 1.0 | 37 ± 7 | 24 ± 16 | 24 ± 5 | 19 ± 2 | 27 ± 10 | 40 ± 14 | 31 ± 15 |
| 31.0 ± 1.5 | 37 ± 9 | 21 ± 5 | 26 ± 3 | 20 ± 3 | 22 ± 12 | 39 ± 14 | 28 ± 9 |
| 43.1 ± 1.6 | 39 ± 9 | 21 ± 3 | 28 ± 4 | 19 ± 4 | 26 ± 12 | 38 ± 10 | 23 ± 6 |
| 84.1 ± 1.1 | 38 ± 9 | 20 ± 3 | 25 ± 3 | 20 ± 3 | 27 ± 13 | 40 ± 10 | 26 ± 7 |

### Running Economy

Footwear bending stiffness did not affect gross aerobic power (p=0.458) (Fig. 7a). Individually, the footwear bending condition that minimized running economy was 13.0 ± 1.0 kN/m for 1 participant, 31.0 ± 1.5 kN/m for 4 participants, 43.1 ± 1.6 kN/m for 4 participants, and 84.1 ± 1.1 kN/m for 6 participants. Also, the footwear bending condition that elicited the worst running economy was 13.0 ± 1.0 kN/m for 5 participants, 31.0 ± 1.5 kN/m for 3 participants, 43.1 ± 1.6 kN/m for 3 participants, and 84.1 ± 1.1 kN/m for 4 participants (Supplementary Fig. 1a-o).

**Figure 7.**
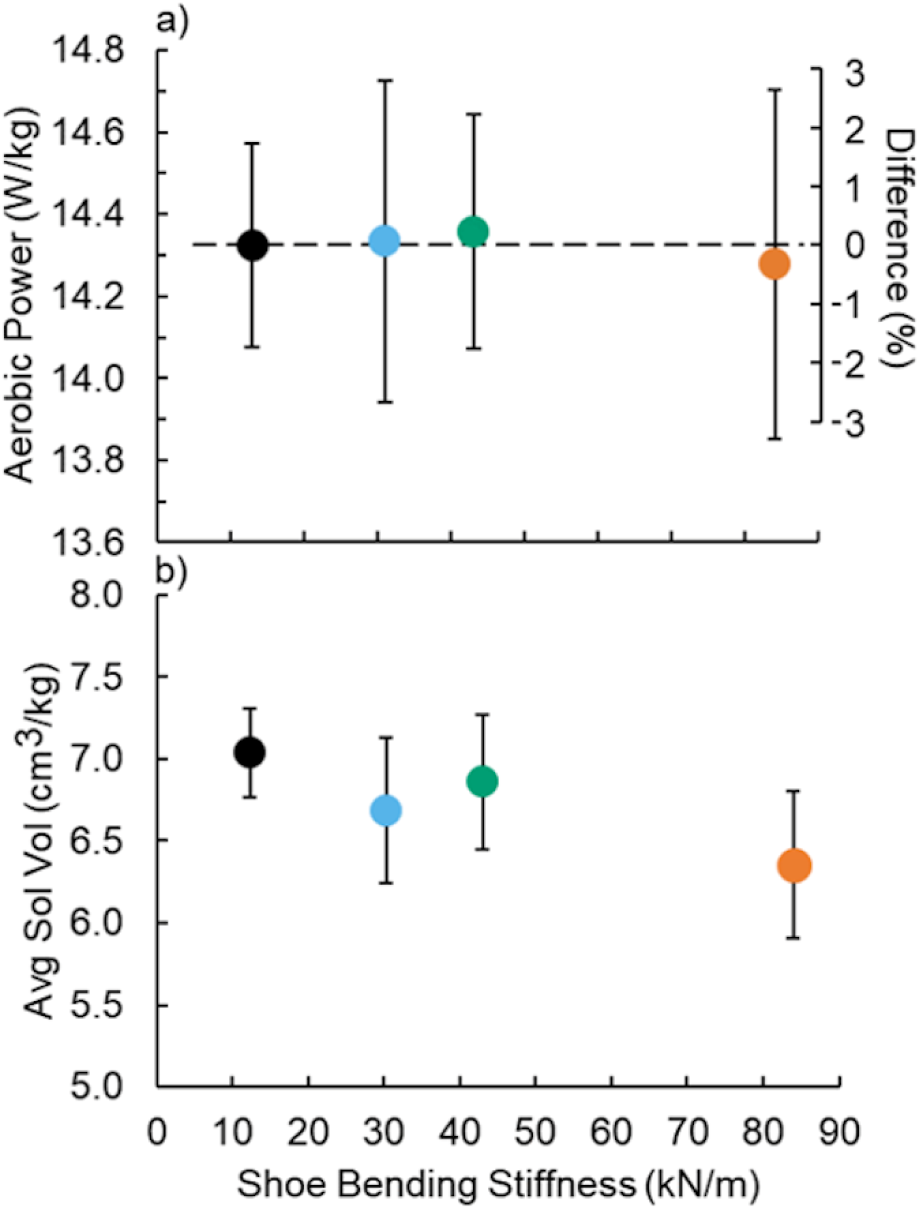
Average (±SE) (a) gross aerobic power and (b) activated soleus (Sol) volume (Vol) per stride. Right axis: Percent difference in the respective variables from the Adidas condition without a carbon fiber plate versus shoe bending stiffness.

## Discussion

Across a 6.4-fold increase in footwear bending stiffness, our participants ran with nearly identical body, limb-joint, and calf muscle mechanics and elicited non-different running economy values. Specifically, footwear bending stiffness did not affect participant GRFs, limb-joint kinematics, or kinetics. Similarly, soleus MT and fascicle dynamics were mostly unchanged across conditions.

Regarding our hypotheses, running in stiffer footwear did not affect soleus force, length, or velocity; thus, we failed to support our initial hypothesis. While no previous study has quantified muscle-level dynamics from athletes running in shoes that varied in bending stiffness, our participant’s unaltered limb-joint dynamics contrasts previous results.^12, 13, 23^ Typically, increased footwear bending stiffness is associated with a more anterior center of pressure and an increased ankle-joint moment.^12, 13, 23^ Still, the only biomechanical difference between our study and the classic investigation that reported that adding carbon fiber plates to shoe soles improve running economy^12^ is that they found an increased maximum ankle moment with the use of stiffer footwear - whereas we did not. Further, while there are likely covariates, athletes running in commercial shoes with versus without curved carbon fiber embedded in their soles exhibited shorter GRF-ankle joint moment arms during ground contact.^35^ Therefore, footwear with increased bending stiffness may not universally increase ankle-joint gear ratio.

The only significant changes that our participants displayed with the use of stiffer footwear were increased ground contact time and reduced stance average and maximum active soleus volume. While considering other factors, longer ground contact times improve running economy by enabling muscles to generate ground force using muscles with more efficient cross-bridge cycling^36^ and reduced active muscle volume improves running economy by requiring fewer energy expending actin-myosin cross-bridges.^34^

Maybe increasing footwear bending stiffness garnered an accumulation of ‘trends’ that yielded significantly less average and maximum active soleus volume per ground contact. Compared to running in the most compliant footwear condition, using the stiffest footwear trended towards a slightly greater duty factor, more plantar flexed ankles, decreased ankle-joint moments, and reduced soleus MT force. Numerically reducing soleus MT force and fascicle pennation angle;^37^ both factors theoretically yield less soleus force and longer soleus fascicles (primarily due to less tendon excursion) with use of the stiffest versus least stiff footwear. Further, the trending direction of active soleus force, length, and velocity with the use of the stiffest versus least stiff footwear likely decreased average and maximum active soleus volume per ground contact.^34^ Again, readers should interpret this paragraph with caution since footwear bending stiffness did not significantly affect any biomechanical measures aside from ground contact time and average and maximum active soleus volume per ground contact.

Despite controlling for shoe mass, adding carbon fiber plates to footwear did not affect running economy nor active soleus volume over the same time-course. Thus, we failed to support our second hypothesis. Additionally, because footwear bending stiffness did not affect the stride-average activation for any of the measured muscles (Table 2 and Fig. 6), none of the respective muscle’s active volume changed across footwear conditions (total muscle volume × relative activation = active muscle volume).^34^ This is the fourth study that failed to replicate the classic investigation^12^ that stated that adding carbon fiber plates to shoe soles improves running economy.^14–16^ Since the classic study, only Oh and Park^13^ reported that adding carbon fiber plates to running shoes to elicit a relative footwear stiffness improves running economy at 2.4 m/s. Moreover, Roy and Stefanyshyn’s original study^12^ reported that participant body mass was inversely correlated with the change in oxygen uptake at their intermediate footwear stiffness condition (38 kN/m) relative to footwear condition that did not have a carbon fiber plate (18 kN/m). Hence, compared to smaller participants, the running economy of larger participants may improve more by adding carbon fiber plates to their shoe soles. In the present study, we found that participant body mass was independent to the change in aerobic power from the most compliant footwear condition in any of the stiffer footwear conditions (all three conditions: p≥0.502). Moreover, due to the implications of muscle dynamics on aerobic power,^34^ we performed post-hoc linear regressions and revealed that the change in aerobic power from the footwear condition that did not have a carbon fiber insole (13.0 ± 1.0 kN/m) was not correlated to the corresponding change in contact time (p=0.135), soleus force generation (p=0.614), soleus velocity (p=0.324), or stride-average soleus muscle volume (p=0.971) (Fig. 8). There was considerable variability regarding how participants altered their running biomechanics with the addition of carbon fiber to their shoe soles (supplementary table 1). Further, there was a potentially spurious correlation between the change in aerobic power and change in soleus length (p=0.040; r=0.311), which indicated that relatively shorter muscle operating lengths are associated with improved running economy - Relatively longer muscles are more economical.^28, 29, 32^ Thus, we did not uncover any muscle-level parameters that correlated with the change in aerobic power when athletes ran in footwear conditions using carbon fiber plates versus without.

**Figure 8.**
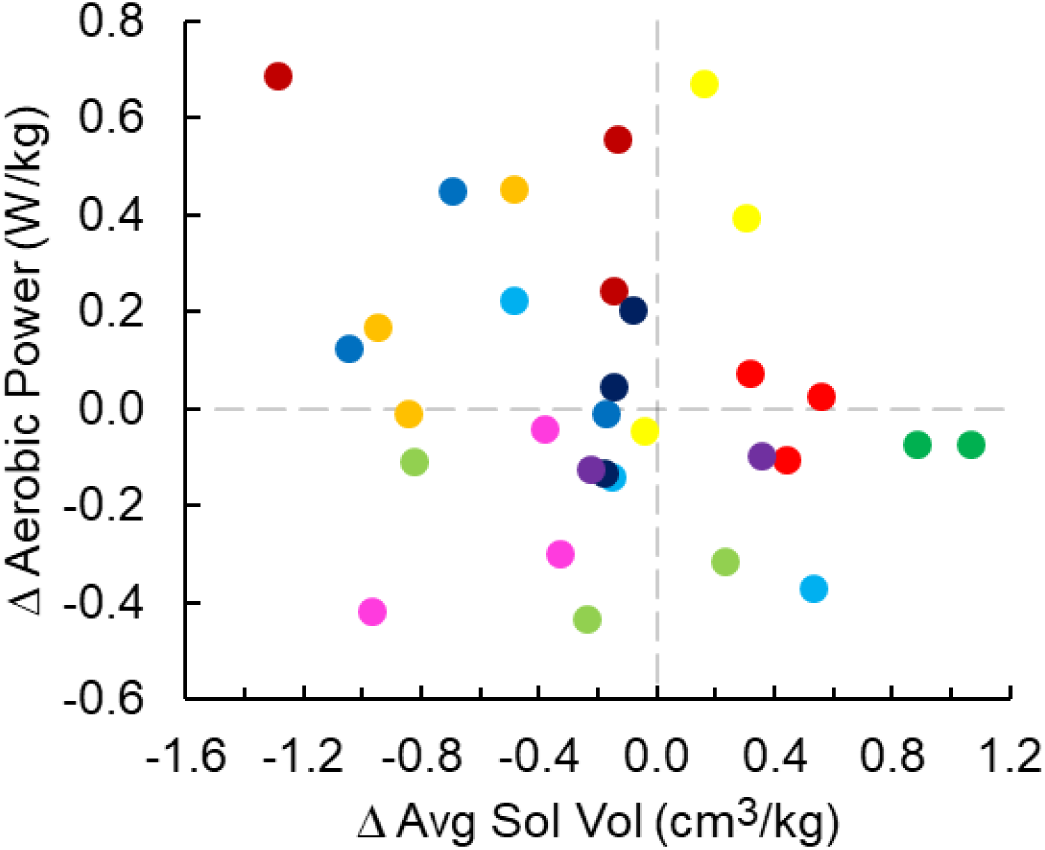
Change in aerobic power versus change in stride-average (Avg) active soleus (Sol) muscle volume (Vol) relative to the Adidas footwear running condition. Each color indicates a different participant. Change in aerobic power was independent of change in Avg Sol Vol (p=0.971).

If footwear bending stiffness does not affect running economy, why does wearing Nike footwear with carbon fiber plates embedded in their midsole (Nike) improve running economy compared to wearing Adidas footwear?^38^ Perhaps Nike’s carbon fiber plate provide the structure necessary for the midsole foam to function. For instance, despite a 264% increased bending stiffness, when athletes run in Nike’s they elicit slightly shorter GRF to ankle-joint moment arms compared to running in Adidas footwear.^35^ This increased footwear bending stiffness and shortened ankle-joint moment arm may be related to Nike’s curved carbon-fiber midsole plates.^38^ Additionally, compared to the Adidas footwear, the respective Nike soles are 8 mm taller (35-62% taller depending on midsole location), the midsole foam is roughly half as stiff (in-series linear stiffness, not bending), and its hysteresis is 11.1% less during vertical loading and unloading.^38^ Because both decreased midsole foam linear stiffness^39–41^ and relative mechanical energy dissipation^42^ in-series to the stance-limb are associated with more economical running, Nike footwear may elicit superior running economy values than Adidas footwear due to their relatively compliant and resilient midsole foam – not increased bending stiffness.

This study has potential limitations. First, our carbon fiber plates were located between the athlete’s sock and the Adidas midsole foam. The lack of cushioning on top of the stiffer carbon fiber plates may have elicited less comfortable footwear compared to the more compliant footwear conditions. However, the vertical GRF magnitude and its average loading rate at 0.05 s (or any preceding timestamp) did not differ between footwear bending conditions (all comparisons p≥0.090), indicating that in-series stiffness or cushioning did not differed or influence our results.^43^ Second, by modeling the soleus as a cylinder,^34^ we likely overestimated participant active soleus volume during running. For example, across footwear conditions, our participant’s estimated maximum active soleus volume was 785 cm^3^, whereas previous studies estimate average total young adult total soleus volume to be 356^44^ to 477^45^ cm.^3^ Yet, because our study focused on the relative changes in active soleus volume with footwear stiffness, this overestimate likely did not influence our results. Third, prior to the experimental trials, each participant performed a five-minute treadmill running habituation trial in the Adidas footwear without a carbon-fiber in-sole. Thus, differences in the habituation time between the footwear bending stiffness conditions may have affected our results. Similarly, both of the previous studies that added carbon fiber plates to shoe soles measured an improved running economy had formal habituation sessions,^12, 13^ whereas we did not. Even though humans adapt their biomechanics in just one step when landing onto terrain with different compliance,^46–48^ perhaps running with carbon fiber insoles requires an extensive habituation period like that of more complicated lower-limb devices (*e.g.* exoskeletons).^49–51^ Additionally, we quantified soleus dynamics and not gastrocnemius dynamics because the soleus is the largest ankle plantar flexor,^45^ the primary muscle that lifts and accelerates the center of mass during locomotion,^52, 53^ likely generates the greatest muscle force of any plantar flexor,^30^ and is often estimated to consume the most metabolic energy of any plantar flexor during running.^30, 54, 55^

## Conclusion

Based on our results, changing footwear bending stiffness hardly changes athlete biomechanics and does not improve running economy. Therefore, if competitive distance runners went back in time, added carbon fiber plates to their footwear, and re-raced, their performance would likely not change.

## Methods

### Participants

Fifteen males participated (Table 1). All participants were apparently free of cardiovascular, orthopedic, and metabolic disorders, and could run 5 km in <25 minutes. Prior to the study, each participant gave informed written consent in accordance with the Georgia Institute of Technology Central Institutional Review Board. During the study. We followed the Georgia Institute of Technology Central Institutional Review Board’s approved protocol and carried out the study in accordance with these approved guidelines and regulations.

### Footwear

We acquired the Adidas Adizero Adios BOOST 2 (Adidas) running shoes in US men’s size 9, 10, 11, and 12. The Adidas are the same shoe model that Dennis Kimetto wore to set a previous marathon (42.2 km) world record (2:02:57 hr:min:s). Next, we fabricated sets of custom carbon fiber in-soles that were 0.8, 1.6, and 3.2 mm thick to fit the Adidas shoes.

We characterized the 3-point bending stiffness of each shoe and in-sole condition following previously described methods.^12, 24, 27^ Briefly, we performed 3-point bending tests by placing each footwear condition in a frame with two supporting bars 80 mm apart. We applied a vertical force to the top of each footwear condition midway between the two supporting bars, approximately where the foot’s metatarsophalangeal joint would be located, using a materials testing machine (Instron, Norwood, MA, USA). We applied force three consecutive times to displace each shoe 10 mm following a 2 N preload (loading rate: 8 mm/s). We calculated footwear 3-point bending stiffness during loading using the average linear slope of the force-displacement data (100 Hz) from the following displacement range: 5 to 9 mm. We also set each athlete’s footwear mass equal to their largest footwear condition, which was the Adidas plus thickest carbon fiber in-sole. For example, the size 9 Adidas shoe is 199 g and its stiffest in-sole was 60 g. Accordingly, we set all size 9 footwear conditions to 259 g by securing mass to the tongue of each shoe.

### Protocol

Each participant completed two experimental sessions. During the first session (aerobic session), participants performed a five-minute standing trial followed by five 5-minute treadmill (Bertec Corporation, Columbus, OH, USA) running trials at 3.5 m/s. Prior to each trial, participants rested for at least five minutes. The first running trial served as habituation to treadmill running in the Adidas footwear (no carbon fiber in-sole). During each subsequent trial, participants ran using a different footwear condition: Adidas as well as Adidas with 0.8, 1.6, and 3.2 mm thick carbon fiber in-soles. We randomized footwear trial order. Each participant’s second session (biomechanics session) occurred at the same time of day and <10 days following their first session. During the second session, participants performed four 2-minute treadmill running trials at 3.5 m/s using the same footwear conditions as the first session in a re-randomized order. We performed separate aerobic and biomechanics sessions to mitigate the potential for technical difficulties to arise by measuring biomechanics over a briefer session than needed for accurate metabolic measurements.

### Aerobic Energy Expenditure

We asked participants to arrive to their aerobic session three-hours post-prandial. Throughout each of the aerobic session’s trials, we utilized open-circuit expired gas analysis (TrueOne 2400, ParvoMedic, Sandy, UT, USA) to record the participant’s rates of oxygen uptake (V̇o_2_) and carbon dioxide production (V̇co_2_). We monitored each participant’s respiratory exchange ratio (RER) throughout each trial to ensure that everyone primarily relied on aerobic metabolism during running; indicated by an RER ≤1.0.^31^ Next, we averaged V̇o_2_ and V̇co_2_ over the last 2-minutes of each trial and used a standard equation^56^ to calculate aerobic power (W). Subsequently, we subtracted the corresponding session’s standing aerobic power (Table 1) from each running trial and divided by participant mass to yield mass-normalized aerobic power (W/kg).

### Biomechanics

Prior to the biomechanics session’s running trials, we placed reflective markers on the left and right side of each athlete’s lower body following a modified Helen Hayes marker set: superficial to the head of the 1^st^ and 5^th^ metatarsal, posterior calcaneus, medial and lateral malleoli, lateral mid-shank, medial and lateral knee-joint center, lateral mid-thigh, greater trochanter, anterior superior iliac crest, posterior superior iliac crest, and superior iliac crest. During the ensuing trials, we recorded vertical and anterior-posterior GRFs (1000 Hz) as well as motion capture (200 Hz) data during the last 30 seconds of each trial. We filtered GRF and center-of-pressure data using a fourth-order low-pass critically damped filter (14 Hz). We filtered motion capture using a fourth-order low-pass Butterworth filter (7 Hz). Using the filtered GRFs, we calculated whole-body stride kinematics (stance and stride time) and GRF parameters (stance average vertical and resultant GRF, as well as mean braking and propulsive horizontal GRFs) with a custom MATLAB script (Mathworks, Natick, MA) that detected periods of ground contact using a 30 N vertical GRF threshold. We categorized each participant as a heel striker or mid/forefoot striker based on visual inspection and whether their vertical GRF trace had an impact peak or not (Table 1). If the participant visually appeared to contact the ground with their heel and displayed a vertical GRF impact peak they were deemed a heel striker.^57^ Participants that did not satisfy these criteria were deemed a mid/forefoot strikers.

We performed inverse dynamics and determined limb joint kinematics (limb joint angles and GRF-to-joint-center moment arms) and kinetics (limb joint moments) (C-motion Inc., Germantown, MD; Mathworks Inc., Natick, MA, USA). Subsequently, we computed each participant’s instantaneous soleus muscle-tendon (MT) moment arm, length, velocity, and force. We used participant anthropometric data and limb-joint angles to calculate the respective soleus MT length,^33^ velocity, and moment arm.^33, 58^ Next, we used each soleus MT moment arm (r) and net ankle-joint moment (M) to calculated soleus MT force (F).

Prior to the biomechanics session’s trials, we secured a linear-array B-mode ultrasound probe (Telemed, Vilnius, Lituania) to the skin superficial of each athlete’s right soleus. Using ultrasonography, we recorded mid-soleus fascicle images (100 Hz) during at least five consecutive strides per trial. We processed the images using a semi-automated tracking software^59^ to determine instantaneous soleus pennation angle and fascicle length. We visually inspected each image to verify the accuracy of the semi-automated tracking software. For semi-automated images that did not accurately track the respective soleus fascicle angle and/or length, we manually redefined the respective fascicle’s parameters. We used soleus MT force and fascicle angle to calculate soleus fascicle force, length, and velocity in congruence with previous studies.^34, 60^ We filtered soleus fascicle angle and length using a fourth-order low-pass Butterworth filter (10 Hz) and took the derivative of fascicle length with respect to time to determine fascicle velocity. Subsequently, we determined relative soleus fascicle length and velocity by deeming that soleus fascicles are at 97% of their resting length at initial ground contact of the Adidas condition^32^ and that their maximum velocity is 436 mm/s,^61^ respectively. Due to technical difficulties, we were unable to compute accurate active soleus volume during 18 of 60 trials; spanning 5 participants.

We recorded surface EMG signals from the biomechanics session’s running trials using the standard procedures of the International Society for Electrophysiology and Kinesiology.^62^ Prior to the first trial, we shaved and lightly abraded the skin superficial to the medial gastrocnemius, soleus, tibialis anterior, vastus medialis, rectus femoris, biceps femoris, and gluteus maximus of each participant’s left leg with electrode preparation gel (NuPrep, Weaver and Co., Aurora, CO). Next, we placed a bipolar surface electrode (Delsys Inc., Natick, MA) over the skin superficial to each respective muscle belly and in the same orientation as the respective muscle fascicle. We recorded EMG signals at 1000 Hz and verified electrode positions and signal quality by visually inspecting the EMG signals while participants contracted the respective muscle. Based on visual inspection and technical difficulties, we removed 97 of 420 potential muscle activation signals due to their poor signal quality; spanning 4 participants. To analyze EMG signals from the running trials, we band-pass filtered the raw EMG signals to retain frequencies between 20 and 450 Hz, full-wave rectified the filtered EMG signals, and then calculated the root mean square of the rectified EMG signals with a 40 ms moving window.^63^ Lastly, we normalized each muscle activation to the average maximum activation of the respective muscle during running in the Adidas condition sans carbon fiber plates.^64^

### Statistics

We performed a linear regression on the footwear’s materials testing force-displacement profile. We performed independent repeated measures ANOVAs to determine whether footwear bending stiffness (independent variable) affects athlete running biomechanics or aerobic power (dependent variables). We performed all statistical tests using R-studio (R-Studio Inc., Boston, USA).

## Acknowledgements

We thank Dr. Frank Hammond and Lucas Tiziani for use of the materials testing machine, in addition to Dr. Young-Hui Chang for the use of his analysis software. We thank PoWeR and EPIC Lab members, namely Lindsey Trejo for assisting data collection and Krishan Bhakta for assisting data analysis. This study was supported by O.N.B.’s National Institute of Health’s, Institute of Aging Fellowship: F32AG063460; P.R.G.’s National Science Foundation Graduate Research Fellowship: DGE-1650044; G.S.S’s National Institute of Health’s, Institute of Aging Award: R0106052017; and G.S.S.’s support from the George W. Woodruff School of Mechanical Engineering.

## Supplementary Files

**Supplementary Figure 1.**
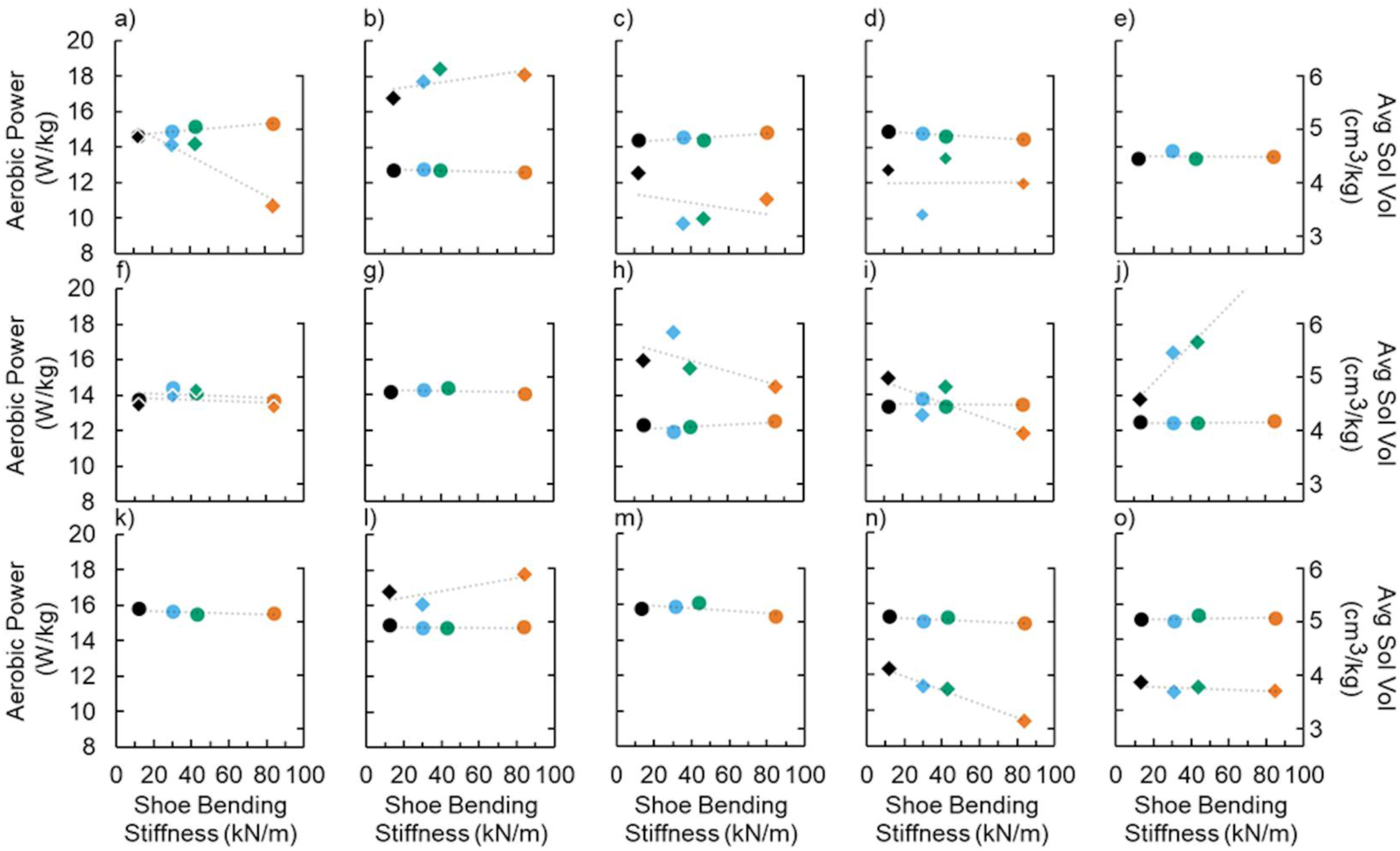
Gross aerobic power (left axis and circles) and stride average soleus (Sol) volume (Vol) versus footwear bending stiffness for each participant.

**Supplementary Table 1.** Number of participants that yielded the lowest and highest value for select biomechanical variables.

| Footwear<br>Bending<br>Stiffness<br>(kN/m) | Ground<br>Contact<br>Time |  | Hip<br>Moment |  | Knee<br>Moment |  | Ankle<br>Moment |  | Soleus<br>Force |  | Soleus<br>Fascicle<br>Length* |  | Soleus<br>Fascicle<br>Velocity* |  |
| --- | --- | --- | --- | --- | --- | --- | --- | --- | --- | --- | --- | --- | --- | --- |
|  | Low | High | Low | High | Low | High | Low | High | Low | High | Low | High | Low | High |
| 13.0 ± 1.0 | 6 | 1 | 7 | 1 | 6 | 3 | 1 | 8 | 4 | 7 | 1 | 3 | 4 | 2 |
| 31.0 ± 1.5 | 3 | 4 | 1 | 4 | 3 | 4 | 6 | 2 | 4 | 2 | 2 | 3 | 1 | 4 |
| 43.1 ± 1.6 | 3 | 3 | 4 | 4 | 1 | 3 | 4 | 1 | 4 | 3 | 5 | 1 | 2 | 4 |
| 84.1 ± 1.1 | 3 | 7 | 3 | 6 | 5 | 5 | 4 | 4 | 3 | 3 | 3 | 4 | 4 | 1 |

\*n=11 participants due to technical difficulties.

